# Seamless Multiscale Modeling of Coagulation Using Dissipative Particle Dynamics

**DOI:** 10.1101/181099

**Authors:** Alireza Yazdani, Zhen Li, Jay D. Humphrey, George Em Karniadakis

## Abstract

We propose a new multiscale framework that seamlessly integrates four key components of blood clotting namely, blood rheology, cell mechanics, coagulation kinetics and transport of species and platelet adhesive dynamics. We use transport dissipative particle dynamics (tDPD), which is the extended form of original DPD, as the base solver, while a coarse-grained representation of blood cell’s membrane accounts for its mechanics. Our results show the dominant effect of blood flow and high Peclet numbers on the reactive transport of the chemical species signifying the importance of membrane bound reactions on the surface of adhered platelets. This new multiscale particle-based methodology helps us probe synergistic mechanisms of thrombus formation, and can open new directions in addressing other biological processes from sub-cellular to macroscopic scales.

## 1 Introduction

Normal hemostasis and pathological thrombosis in a flowing bloodstream can occur on damaged tissues in the normal circulation, especially over exposure to collagen and surfaces of artificial internal organs. The clot is predominantly composed of platelets, which are nondeformable ellipsoid shape blood cells in their unactivated state circulating in blood with a concentration of 150, 000 — 400, 000 *mm*^−3^. Thrombus formation and growth at a site of injury on a blood vessel wall is a complex biological process, involving a number of multiscale simultaneous processes including: multiple chemical reactions in the coagulation cascade, species transport, cell mechanics, platelets adhesion and blood flow. Thus, numerous computational models have been developed to address the underlying hemostatic processes as a whole or individually (see reviews in [22, 24]) providing useful information on the mechanism of clotting and its growth despite making assumptions and simplifications in the models. Incorporating all these processes that occur at different spatio-temporal scales, however, remains a rather challenging task. For example, platelet-wall and platelet-platelet interactions through receptor-ligand bindings occur at a sub-cellular nanoscale, whereas the blood flow dynamics in the vessel around the developing thrombus is described as a macroscpoic process from several micrometers to millimeters.

There are a few multiscale computational models attempting to address the whole clotting process or some of its major subprocesses *e.g.*, by Xu et *al.* [25], Wu et *al.* [23], Tosenberger *et al.* [21], Flamm *et al.* [7] and Fogelson and Guy [8]. Xu *et al.* outlined a multiscale model that included macroscale dynamics of the blood flow using the continuum Navier-Stokes (NS) equations, and microscale interactions between platelets and the wall using a stochastic discrete cellular Potts model. The enzymatic reactions of the coagulation pathway at the injured wall in plasma, as well as on the platelet’s surface were incorporated through coupling advection-diffusion-reaction (ADR) equations to blood flow. The numerical simulations, however, remained two-dimensional due to the significant cost of solving ADR equations. Flamm *et al.* presented a patient-specific multiscale model, in which patients' data were used to train a neural network model for platelet’s calcium signaling. The background blood flow was solved using lattice Boltzmann method (LBM), whereas the concentration fields for the important agonists namely, adenosine diphosphate (ADP) and thromboxane A2 (TxA2), were solved using finite elements. A stochastic lattice kinetic Monte Carlo method was used for microscopic adhesive kinetics. Similar to the work by Xu et al., this study was also limited to 2D microchannels, where platelets were treated as circular particles. Wu *et al.* introduced a three-dimensional bead-spring representation of platelet’s membrane, whereas they used LBM to model blood flow, and the immersed boundary method to couple platelet and blood interactions. Tosenberger *et al.* [21] developed a hybrid model where dissipative particle dynamics (DPD) is used to model plasma flow and platelets, while the regulatory network of plasma coagulation is described by a system of partial differential equations governing the ADR equations. Fogelson and Guy [8] used a continuum Eulerian-Lagrangian approach, where the NS equations were solved along with the ADR equations for the transport of species, whereas an immersed boundary method was used to couple platelet (treated as rigid spherical particles) interactions with blood.

As mentioned above, there is a high level of complexity and heterogeneity in the current proposed multiscale models, limiting their ability in dealing with an involved three-dimensional and physiologic clotting process. Our goal in this work is to introduce a multiscale numerical framework that seamlessly unifies and integrates different subprocesses within the clotting process. Therefore, we are able to reduce the degree of heterogeneity in the previous models, and increase the efficiency of simulations by cutting the unnecessary communication overhead between different solvers.

To this end, we employ dissipative particle dynamics, a mesoscopic particle-based hydrodynamics approach [3,9], to model plasma and suspending cells in blood. DPD has been shown to be an effective particle-based method to study cell motions and interactions in suspensions such as blood [6]. The advantage is the seamless integration of hydrodynamics and cell mechanics in a single framework. Further, we utilize our recently developed transport DPD (tDPD) [15] to address chemical transport in the blood coagulation process, which drastically reduces the cost of solving ADR equations. The details of the numerical methodology is given in Section 2.

## 2 Multiscale numerical methodology

The foundation of our multiscale model is the DPD numerical framework upon which other modules are established. The multiscale model includes four main modules: the hemodynamics, coagulation kinetics and transport of species, blood cell mechanics and platelet adhesive dynamics. DPD can provide the correct hydrodynamic behavior of fluids at the mesoscale, and it has been successfully applied to study blood, where a coarse-grained representation of cell membrane is used for both RBCs [5, 18] and platelets [26]. The DPD representation of red cells in blood was extensively used and validated in the previous studies for suspensions of both healthy [6] and disease cells [14] (*e.g.*, malaria and sickle cells). The following sections describe each module in more detail.

### 2.1 Blood rheology

In the standard DPD method, the pairwise forces consist of three components: (i) conservative force 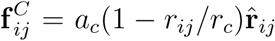; (ii) dissipative force, 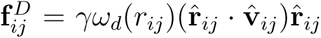; and (iii) random force, 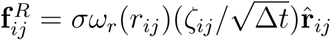. Hence, the total force on particle *i* is given by 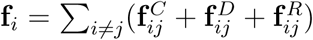, where the sum acts over all particles *j* within a cut-off radius *r*_c_, and *a*_c_, γ, σ are the conservative, dissipative, random coefficients, respectively, *r*_*ij*_ is the distance with the corresponding unit vector 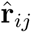, 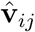 is the unit vector for the relative velocity, 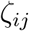 is a Gaussian random number with zero mean and unit variance, and Δt is the simulation timestep size. The parameters γ and σ and the weight functions are coupled through the fluctuation-dissipation theorem and are related by 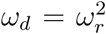 and 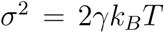, where *k*_*B*_ is the Boltzmann constant and *T* is the temperature of the system. The weight function *ω_τ_(r_ij_)* is given by

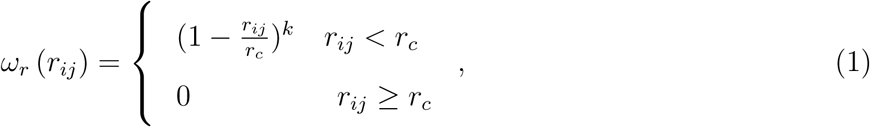

with *k* = 1 in the original DPD form, whereas *k* = 0.2 is used in this study to increase the plasma viscosity. Table 1 presents the DPD parameters used for the fluid (representing plasma) particles throughout this study.

**Table 1:** DPD fluid parameters used in this study: *n* is the fluid’s number density, a_c_ is the conservative force coefficient, γ is the dissipative force coefficient, k is the weight function exponent, and *η* is the fluid’s dynamic viscosity. In all simulations, we set the particle mass m =1, and the thermal energy k_B_T = 0.1 in DPD units.

| $n$ | $r_c$ | $a_c$ | $\gamma$ | $k$ | $\eta$ |
| --- | --- | --- | --- | --- | --- |
| 4.0 | 1.58 | 5.0 | 20.0 | 0.2 | 117.6 |

### 2.2 Coagulation kinetics and transport

Transport of chemical species in the coagulation cascade is modeled by tDPD, which is developed as an extension of the classic DPD framework [15] with extra variables for describing concentration fields. Therefore, equations of motion and concentration for each particle of mass mm are written by (dr_*i*_ = v_*i*_ dt; dv_*i*_ = f_*i*_/*m*_*i*_dt; d*C*_*i*_ = *Q*_*i*_/*m*_*i*_ dt) and integrated using a velocity-Verlet algorithm, where *C*_*i*_ represents the concentration of a specie per unit mass carried by a particle, and *Q*_*i*_ = 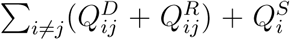 is the corresponding concentration flux. We note that *C*_*i*_ can be a vector C_*i*_ containing *N* components *i.e.* {C_1_, C_2_,C_*N*_}_*i*_ when *N* chemical species are considered. In the tDPD model, the total concentration flux accounts for the Fickian flux 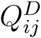 and random flux 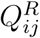, which are given by

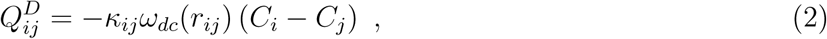

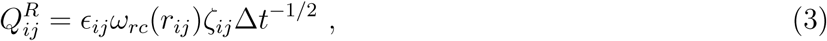

where *κ_ij_* and *ε_ij_* determine the strength of the Fickian and random fluxes, and 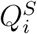 represents the source term for the concentration due to chemical reactions. Further, the fluctuation-dissipation theorem is employed to relate the random terms to the dissipative terms *i.e.*, 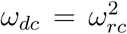, and the contribution of the random flux 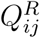 to the total flux is negligible [15].

The effective diffusion coefficient *D* of a tDPD system is a composed of the random diffusion *D*^*ξ*^ and the Fickian diffusion D^F^, and is a linear function of Fickian friction *κ*. Further, *D* has a minimum of *D*_*min*_ = *D*^*ξ*^ at *κ* = 0. Therefore, the maximum Schmidt number (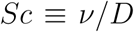) that a tDPD system can achieve is determined by *Sc*(*κ* = 0) = *v*/*D*_*min*_. The DPD parameters considered for plasma are given in Table 1, which lead to plasma kinematic viscosity *ν* = 29.4. In addition, the weight function for chemical transport is taken as ω_*rc*_(*r*) = (1 − *r*/*r*_*cc*_), where the concentration cut-off radius is *r*_*cc*_ = 1.0. Computations for the coefficient of self diffusion based on this system will result in *D*_*min*_ = 3.55 × 10^−4^. Therefore, the diffusion coefficients for all species can be evaluated as *D*_*i*_ = 3.55 × 10^−4^ + 0.0682 *κ*_*i*_, where all cross-diffusions are assumed to be negligible (*i.e.*, *κ_ij_* = 0 for *i* ≠ *j*).

### 2.3 Cell mechanics

In this study we use a coarse-grained representation of the cell membrane structure (lipid bilayer + cytoskeleton), which was introduced in the work of Fedosov *et al.* [5] for red blood cells (RBCs) and extended for platelets in [26]. The membrane is defined as a set of *N_v_* DPD particles with Cartesian coordinates *x*_*i*_, *i* ∈ 1,…, *N*_*v*_ in a two-dimensional triangulated network created by connecting the particles with wormlike chain (WLC) bonds. The free energy of the system is given by

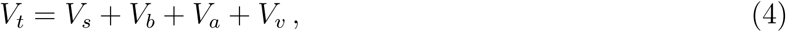

where *V*_*s*_ is the stored elastic energy associated with the bonds, *V*_*b*_ is the bending energy, and *V*_*a*_ and *V*_*v*_ are the energies due to cell surface area and volume constraints, respectively. RBCs are extremely deformable under shear, whereas platelets in their unactivated form are nearly rigid. We parametrize the RBC membrane such that we can reproduce the optical tweezer stretching tests [2]. In the case of platelets, we choose shear modulus and bending rigidity sufficiently large to ensure its rigid behavior. Platelets become more deformable upon activation through reordering the actin network in their membrane, which is accompanied by the release of their content to plasma [27]. This process is not included in our model for simplicity. Further, the plasma-cell interactions contribute to the overall rheology of blood. These interactions are accounted for through viscous friction using the dissipative and random DPD forces, where the repulsive force coefficient for the coupling interactions is set to zero. The strength of the dissipative force is computed such that no-slip condition on the cell surface is enforced. Further details on the RBC and platelet membrane mechanics and plasma-cell interactions can be found in [5, 26].

### 2.4 Platelet adhesive dynamics (PAD)

The PAD model describes the adhesive dynamics of receptors on the platelet membrane binding to their ligands. The cell adhesive dynamics algorithm was first formulated in the work of Hammer and Apte [10] and extended to platelets by Mody and King [16]. This model utilizes the Monte Carlo method to determine each bond formation/dissociation event based on specific receptor-ligand binding kinetics. We estimate the probability of bond formation *P*_*f*_ and probability of bond breakage *P*_*r*_ using the following equations

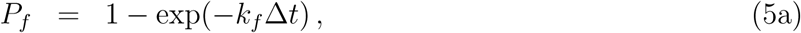

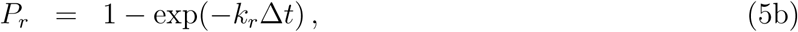

where *k*_*f*_ and *k*_*r*_ are the rates of formation and dissociation, respectively.

Each platelet has approximately 10,688 GPIbα receptors uniformly distributed on its surface [20]. We assume that circulating platelets can form bonds with A1 domains of immobilized vWF multimers on the vessel wall injury, thus excluding the effect of other adhesion mechanisms in our study. Following Mody and King [16], the force-dependent *k*_*f*_ and *k*_*r*_ are evaluated by

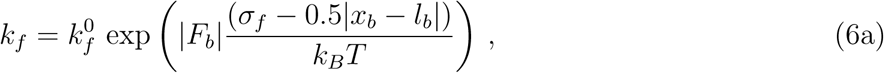

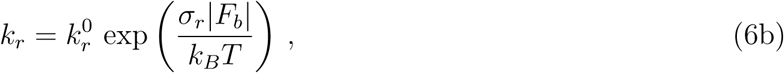

where 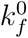 and 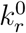 are the intrinsic association and dissociation rates, respectively, and σ_f_ and σ_r_ are the corresponding reactive compliances. Assuming a harmonic GPIbα-vWF bond with stiffness *κ*_*b*_, the force *F*_*b*_ in the bond can be calculated as *F*_*b*_ = − *κ*_*b*_(*x*_*b*_ − *l_b_*), where *x*_*b*_ is the receptor-ligand distance and *l*_*b*_ is the equilibrium bond length. The parameters in Table 2 for the forward and reverse rates are taken from Kim *et al.* [12], where it is shown that GPIbα-vWF bonds behave like “flex” bonds switching to a faster forward rate for bond forces above *F*_*b*_ = 10 *pN*.

**Table 2:** Values for bond formation and breakage kinetics in physical and DPD units. DPD values are estimated based on [*L*] = 1.0 × 10^−6^*m*, [*t*] = 2.27 × 10^−4^ *s*, and [*F*] = 4.5 × 10^−14^ *N*.

| Parameter (unit) | Physical Value | DPD Value | Reference |
| --- | --- | --- | --- |
| $k_f^0$ ($s^{-1}$) | 40.0 | $9.1 \times 10^{-3}$ | [12] |
| $k_r^0$ ($s^{-1}$) | 0.0022 | $5.0 \times 10^{-7}$ | [12] |
| $\sigma_f$ (nm) | 1.8 | $1.8 \times 10^{-3}$ | [12] |
| $\sigma_r$ (nm) | 1.62 | $1.62 \times 10^{-3}$ | [12] |
| $\kappa_b$ (pN/nm) | 10 | $2.222 \times 10^5$ | [16] |
| $l_b$ (nm) | 128 | 0.128 | [16] |

### 2.5 Problem setup

In this work, we setup a 3D straight microchannel with the size of 110 × 20 × 20 *μm*^3^ filled with whole blood at 25% hematocrit, and 500,000 *mm*^−3^ platelet density, where the channel is periodic in *x* and *z* directions. A snapshot of the blood in the channel is shown in Figure 1(a,b). RBCs in their resting form are biconcave with diameter of 7.82 *μm*, whereas platelets are oblate spheroids with diameter of 2 and aspect ratio of 0.3 [19]. The platelet density is taken somewhat larger than the physiologic value, and all platelets are initially placed on the lower side of the channel to accelerate margination and the adhesion process. The coarse-grained RBC membrane is described by 362 DPD particles, whereas the platelet membrane contains 42 DPD particles.

**Figure 1:**
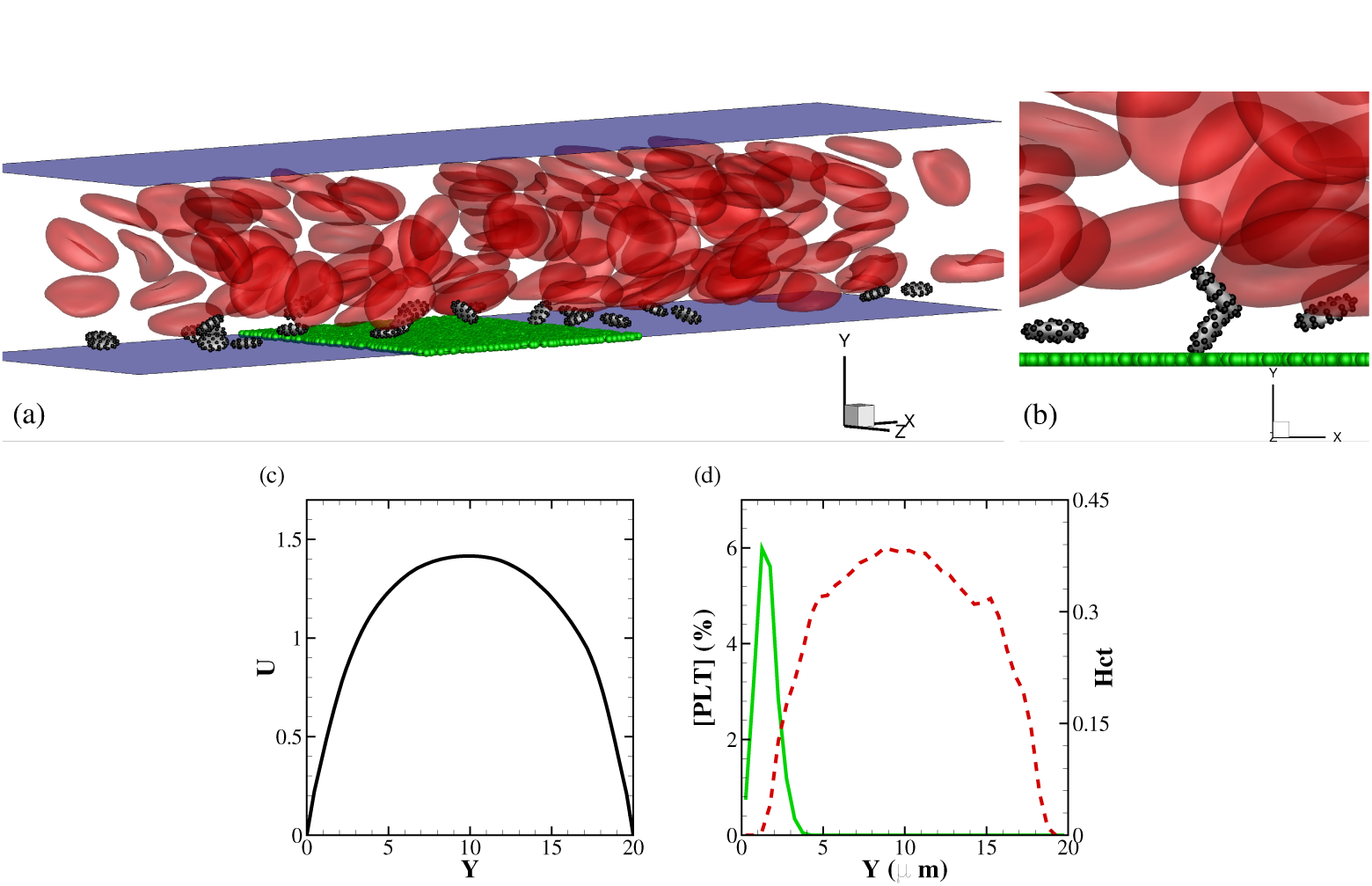
(a) 3D presentation of the channel containing blood at 25% hematocrit. The blue planes represent the channel walls, whereas the green particles are attached to the lower wall and represent the vWF adhesive sites. (b) Closer view at the lower wall showing the particles on platelet’s surface that can bind to ligands upon platelet’s activation. (c) Blood flow velocity profile across the channel height (dimensions are in DPD units). (d) Average cell distribution profiles across the channel height for platelet concentration (solid line and left axis), and blood hematocrit (dashed line and right axis).

To drive the blood flow in a straight channel we apply a constant body force to each DPD particle. This will yield a plug-like profile for blood. The characteristic shear rate is defined as 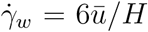, which corresponds to the wall shear rate of a parabolic Poiseuille flow with the same mean velocity *ū*. The Reynolds number is defined based on the channel width H = 20 *μm*, which is *Re* = *ūH*/*v*. The flow condition resembles blood flow in arterioles.

In Figure 1 (c), we plot the plug-like velocity profile for the blood flow from which the mean blood velocity *ū* can be computed. Consequently, the flow *Re* and the wall shear rate 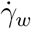 are estimated to be 0.045 and 1400 s^−1^, respectively. Further, the Peclet number, which is the measure of rate of advection to diffusion in transport processes is defined by *Pe* = *Re Sc*, which is ≈ 696 for thrombin (assuming *D*/_*IIa*_ = 6.47 × 10^−7^ *cm*^2^/*s*) in our simulations. We also plot the average distribution of the red cells and platelets across the channel height in Figure 1(d). We observe the presence of a cell-free layer adjacent to the walls with the thickness of ≈ 3 *μm*, where the concentration of platelets is maximum.

Regarding the coagulation cascade, we follow the mathematical model of Anand *et al.* [1], which contains a set of twenty-three coupled ADR equations for describing the evolution of 25 biological reactants involved in both intrinsic and extrinsic pathways of blood coagulation, and the fibrinolysis processes. This model was implemented and tested in our previous work [15]. The ADR equations are in the form of

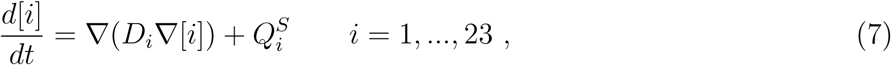

where [*i*] denotes the concentration of *i*^*th*^ reactant and *D*_*i*_ is the corresponding diffusion coefficient. 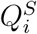 are the nonlinear source terms representing the production or consumption of [*i*] due to the enzymatic reactions. Here, we focus only on tissue factor (TF) or extrinsic pathway to isolate and analyze the effect of TF-VIIa complex on the rate of thrombin production. The details of the bulk and surface chemical reactions along with their relevant constants are given in Li *et al.* [15]. The initial concentration of Fibrinogen (I), zymogens (II, V, VIII, IX, X, XI), inhibitors (ATIII, TFPI), and protein C (PC) are determined based on normal human physiological values, whereas concentrations of the enzymes (IIa, Va, VIIIa, IXa, Xa, XIa), fibrin and APC in plasma are initially zero [1]. Two surface reactions on the wall at the site of injury will generate enzymes IXa and Xa, which in turn, initiates the extrinsic coagulation cascade. Surface-bound TF-VIIa complex is required in the surface reactions and its concentration is taken as 200 *fM* in our study [1]. We also consider two different concentration levels of 50 and 400 *fM* for sensitivity analysis.

The wall boundaries in the simulation domain are treated as pseudo-planes similar to the work of Lei *et al.* [13]. Specifically, particles within one cut-off distance from the wall are reflected in a bounce-forward scheme. Further, in order to impose the no-slip condition, a predefined force that compensates for the missing particles on the opposite side of the wall is applied to the particles both in wall normal and tangential directions. Following Li *et al.* [15], in order to impose the Dirichlet and Neumann boundary conditions for the concentrations, we apply a predefined flux to the particles within one cut-off radius of the wall.

In order to relate the DPD parameters with the physical values, we need to first define length and time scales. The RBC membrane shear modulus imposes the time scale for the DPD system, which follows

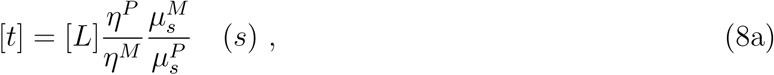

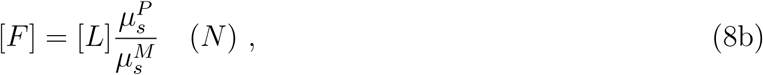

where *μ_s_* is the RBC membrane shear modulus, *η* is the plasma viscosity, and superscripts *M* and *P* denote the model (DPD) and physical units, respectively. The length scale is taken as [*L*] = 1 × 10^−6^ *m*, whereas the time scale is evaluated by Eq. (8) [*t*] = 2.27 × 10^−4^ s (using membrane shear modulus 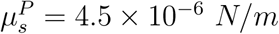 and plasma viscosity *η*^*P*^ = 1.2 × 10^−3^ *Pas*). Further, chemical concentrations are scaled by [C*0*] = 1.0*nM* · [*L*]^3^/*n*, where *n* is the DPD particles density.

We also note that the physiologic timescale of blood coagulation dynamics is in the order of minutes, hence making the long-time tDPD computation expensive and almost impractical. Therefore, in our numerical simulations, the reaction kinetics have to be accelerated by a factor of *O*(100). However, as we keep the transport timescale defined by the Peclet number unchanged (the compounds diffusion coefficients and flow velocity remains the same as the original process), any acceleration in chemical reaction timescales may induce a discrepancy in the rate of thrombin generation from the original non-accelerated process. To further asses this deviation, we present an analysis for three different levels of acceleration in the following section.

## 3 Results

We first plot a snapshot of the whole blood simulation in the channel at time *t* = 34 *s* (see Figure 2), where the concentration boundary layer has been fully developed (shown by contours of thrombin concentration [*IIa*] in the *x* − *y* plane) and few platelets are adhered to the site of injury (shown by green particles). We set a threshold value of 1 *nM* for thrombin-mediated platelet activation [17] for which the platelets in the boundary layer become instantaneously activated and ready to form bonds with the particles representing vWF ligands on the site of injury. The thickness of thrombin boundary layer is approximately 2 *μm*, which is reasonable under the current flow conditions and Peclet number.

**Figure 2:**
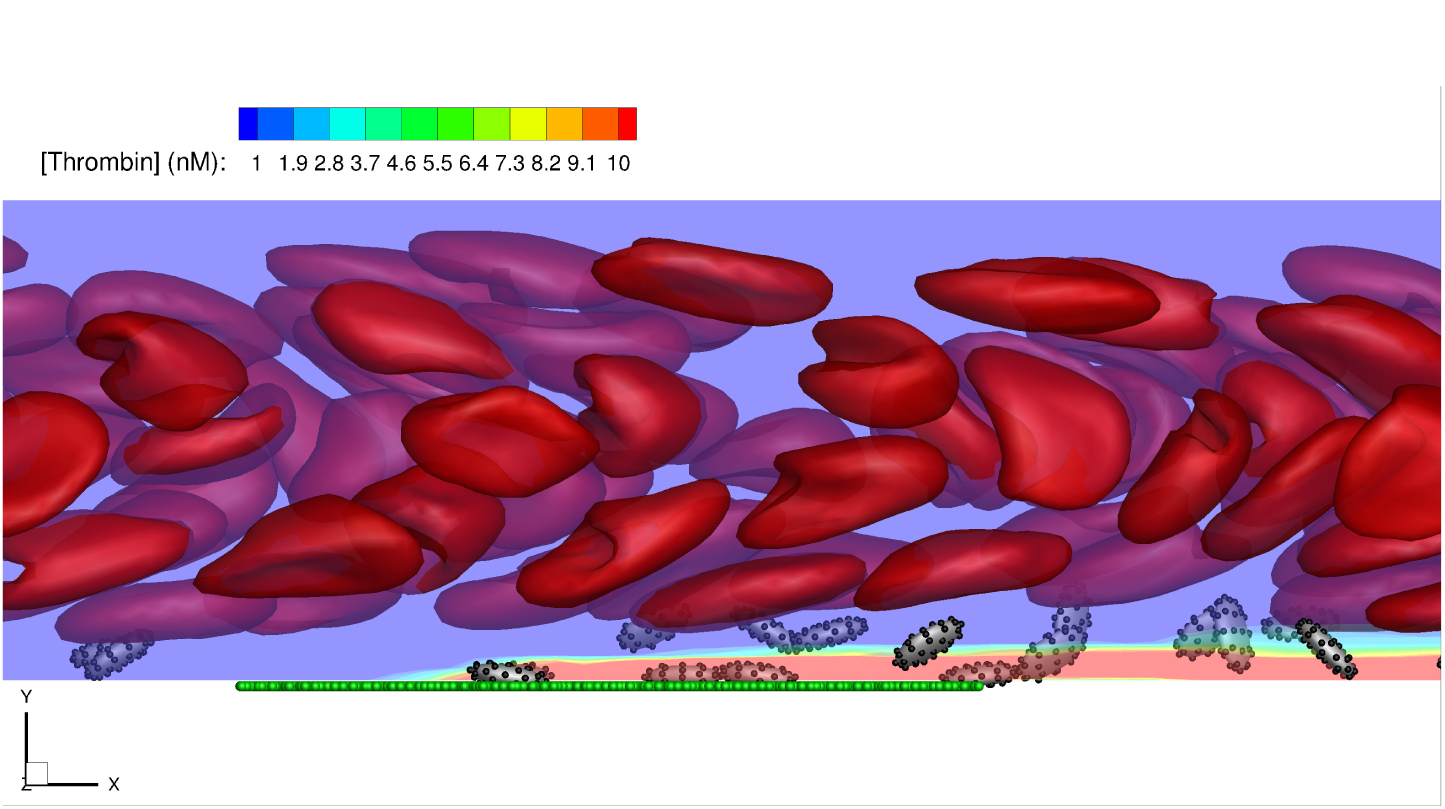
Snapshot of the DPD numerical result showing red blood cells, platelets, vWF ligands (green particles on the lower wall), and contours of thrombin concentration. Also shown, few platelets that are adhered on the injured area (a video for the full simulation is given in the Supplementary Material).

Figure 3(a) presents a closer look at the time-evolution of thrombin concentration at three different axial positions, *X* = 40, 50, 60 *μm*. Here, the reaction rates for the bulk reactions are accelerated 100 times, while the surface reaction rates are not accelerated. The results show no generation of thrombin up to *X* = 50 *μm*, at which thrombin production is moderate. A more rapid burst of thrombin followed by a decrease in the concentration is observed at *X* = 60 *μm*. This observation shows the strong effect of flow rate on reactive transport of chemical species in the coagulation cascade. The slow kinetic rates delay thrombin production toward the end of injury site as blood flow tends to transport the activated compounds IXa and Xa downstream. This result is more pronounced in Figure 3(b) as we use smaller acceleration factors for the kinetic rates (*i.e*., 25 and 10 times faster the measured values). We observe much less production of thrombin for most part of the injury site, whereas a significant burst of thrombin occurs toward the end of site of injury due to accumulation of transported activated compounds in that region. Taken together, these results underline the important role of blood flow in the transport of species away or into the site of injury. More importantly, they imply that membrane-bound kinetics, which may occur on the surface of adhered platelets (not included in this study), are essential to thrombin production throughout the propagation phase. We replot concentrations of Figure 3(b) with respect to non-accelerated time, where the original timesacle [*t*] = 2.27 × 10^−4^ *s* has been used. Interestingly, we observe that changes in the kinetic rates affect the magnitude of thrombin concentration whereas the rate of concentration growth is almost unaffected. This result also indicates the stronger rate of advective transport compared to the rate of chemical reactions under such flow conditions.

**Figure 3:**
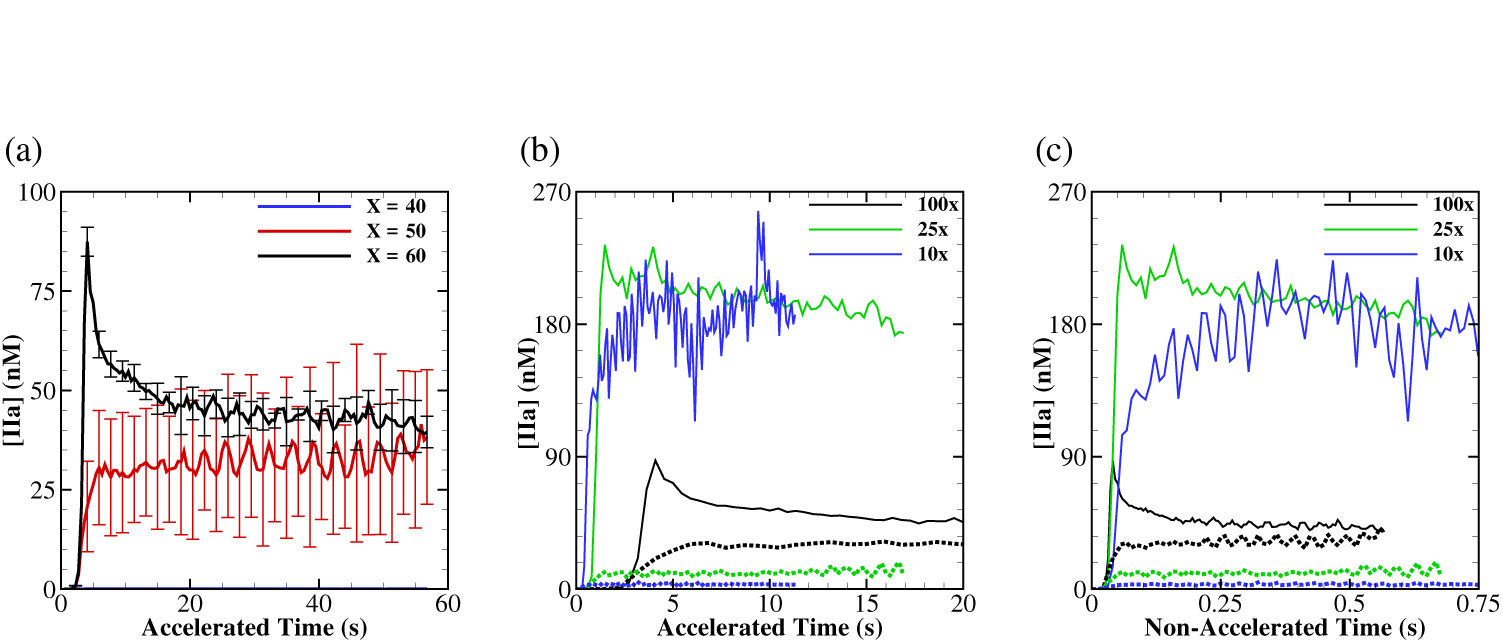
(a) Evolution of thrombin concentration at three axial positions on the site of injury for 100 times accelerated bulk kinetic rates vs. accelerated time (*i.e.*, 100 × [*t*] = 2.27 × 10^−2^ *s*). Comparison of thrombin concentration for three levels of acceleration plotted against: (b) accelerated time; (c) non-accelerated time (**i.e.**, [*t*] = 2.27 × 10^−4^ *s*). Dotted and bold lines are extracted for *X* = 50 *μm* and *X* = 60 *μm*, respectively. The surface concentration of TF-VIIa complex is taken as 200 *f M* for all cases.

We plot the total number of bonds between activated platelets and the ligands in Figure 4(a) with respect to time for five simulations under the same flow conditions. In addition, Figure 4(b) shows the number of adhered platelets with time. After an initial time lag of approximately 5 seconds, the number of bonds and adhered platelets both start to rise sharply, and eventually reach a plateau as platelets cover most of the injured area. The *in vivo* experimental measurements of Falati *et al.* [4] based on fluorescence microscopy in mice arterioles also show similar trends of platelet aggregation as thrombus develops. More specifically, they observe a time lag in the initiation of platelet aggregation followed by the increase in fluorescence intensity (*i.e.*, number of adhered platelets) that also reaches a plateau. Although the numerical growth curves are qualitatively similar to experimental observations, we should highlight that in our simulations platelets only interact with the wall, which leads to formation of a monolayer. Platelet-platelet interactions are facilitated through their membrane receptors and ligands, and can easily be included in the numerical model.

**Figure 4:**
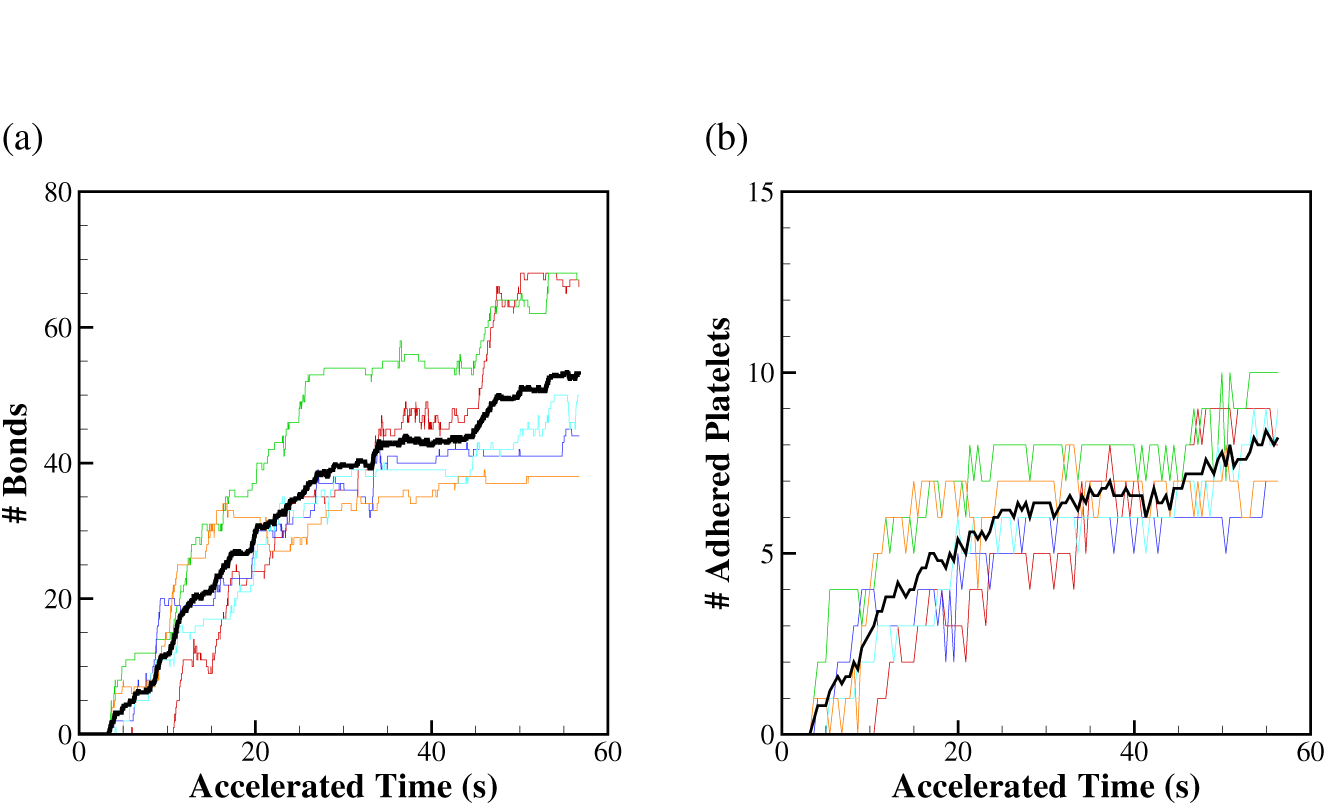
(a) Total number of bonds formed between adhered platelets and vWF ligands as a function of time for 5 ensemble runs (light colored lines) including their mean (black bold line). (b) Number of platelets adhered to the site of injury vs. time for 5 different runs along with their mean curve. Here the surface concentration of TF-VIIa complex is taken as 200 *f M*, and reaction rates are accelerated 100 times.

We also present the sensitivity of the results to several other important parameters *i.e.*, to the boundary concentration level of TF-VIIa complex and platelet’s adhesive parameters. Figure 5(a) shows the effect of TF-VIIa concentration at the site of injury on the number of bond formation. As can be seen here lower concentration of this complex at around 50 fM is almost ineffective in the platelet aggregation process, whereas higher levels of this complex by several factors will not change aggregation rate dramatically. In addition, we tested the effect of binding site density on the platelet surface as well as the bond intrinsic dissociation rate on the number of bond formations. It should be emphasized that, while there is a significant number of GPIba receptors on platelet’s surface, not all of them are able to form bonds. These receptors are normally distributed uniformly on the platelet’s surface. Platelets, on the other hand, move very close to the wall when they perform a tumbling motion exposing only a very small region of their surface to vWF ligands on the wall. Here, we consider two different scenarios: first we increase the intrinsic bond dissociation rate by an order of magnitude to 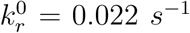; second, we lower the number of binding sites on platelet’s surface by a factor of 3 compared to its physiologic value; the latter can be associated to some glycoprotein receptor deficiencies such as Bernard-Soulier syndrome that is characterized by loss-of-function mutations in GPIba [11]. The results are shown in Figure 5(b) and compared with the control. As expected the higher rate of dissociation reduces the number of bond formations and aggregated platelets. The reduction in the number of receptors on platelet’s surface adversely affects the adhesive dynamics causing a significantly less number of bond formations.

**Figure 5:**
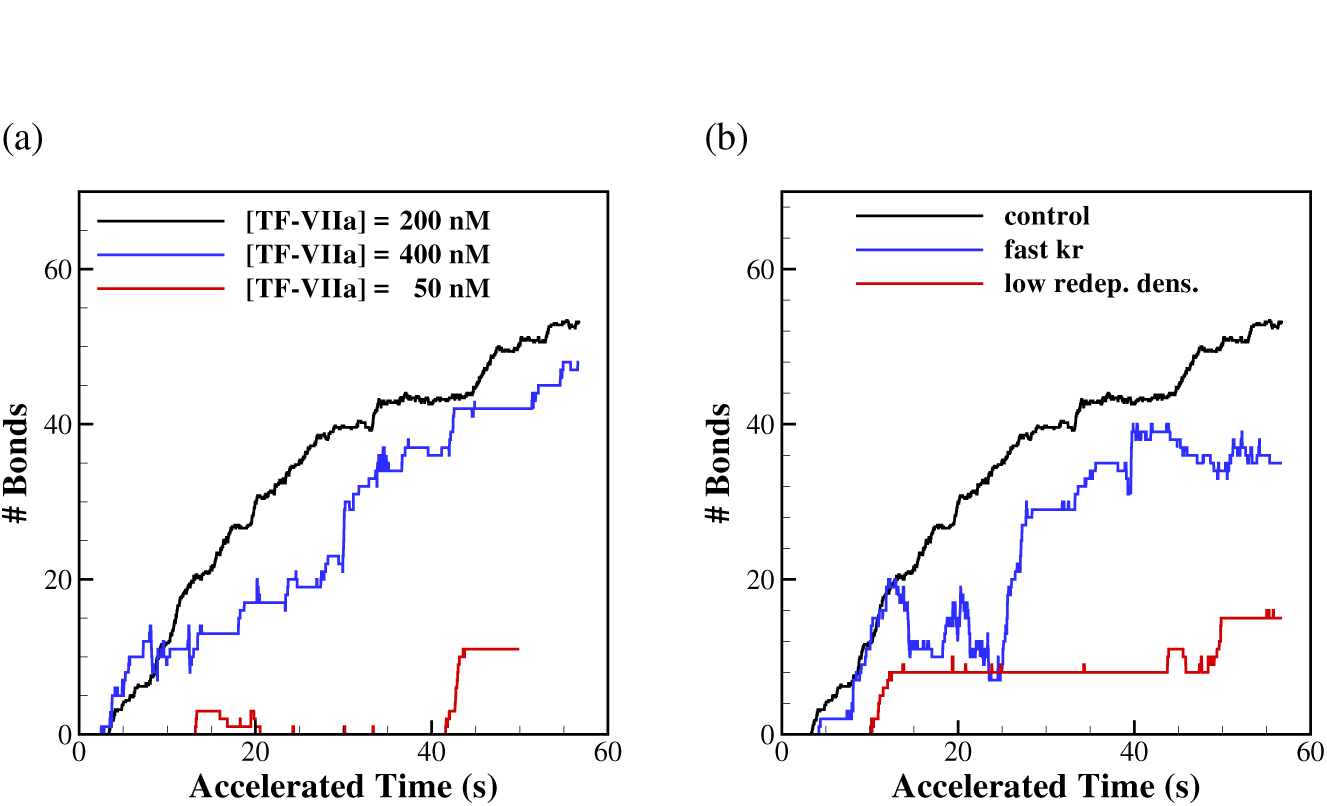
The effect of TF-VIIa concentration and platelet adhesive parameters on the number of bond formations: (a) total number of bonds formed between adhered platelets and vWF ligands as a function of time for three levels of TF-VIIa concentration; (b) total number of bonds formed for increased 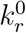 and reduced number of binding sites. Control is for 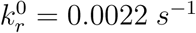 and 10,688 receptors on platelet’s surface. Here the reaction rates are accelerated 100 times for all cases.

## 4 Conclusion

We presented a multiscale numerical framework based on transport DPD method to address the complicated biophysical processes involved in blood clotting. The numerical model seamlessly integrates the relevant sub-processes *e.g.*, sub-cellular platelet adhesive dynamics, mechanics of blood cells, biochemistry of coagulation cascade and blood rheology.

To test the numerical model and perform sensitivity analyses for different model parameters, we used a straight channel as a simplified geometry for blood clotting simulations. Results show good qualitative agreement with experiments for the flow conditions in arterioles. We believe such a numerical framework will be able to effectively address different multiscale biological processes and can be extended to more complicated systems and geometries as well.

## 5 Acknowledgement

This work is funded by the NIH Grant No. U01HL116323. AY would like to thank the computational resources provided by NSF-XSEDE (SDSC Comet and TACC Stampede) through the award No. TGDMS140007. ZL would like to thank the computational support and resources from DOE/ANL and DOE/ORNL through an INCITE award.

